# Dorsal hippocampal oxytocin receptor regulates adult peer bonding in rats

**DOI:** 10.1101/2023.12.04.570022

**Authors:** Yufei Hu, Wensi Li, Yinji Zhao, Yuying Liu, Wenyu Sun, Yi Yan, Laixin Liu, Bowen Deng, Pu Fan

## Abstract

Healthy social relationships are beneficial whereas their breakdown is often linked to psychiatric disorders. Parental care and bonding between sexual partners have been well studied both at the level of behavioral analysis and underlying neuronal mechanisms. By contrast, little is known about the neural and molecular basis of peer bonding, defined as social bonds formed between unrelated individuals of the same sex, due to the lack of a suitable experimental paradigm. We found that adult Sprague Dawley (SD) rats of the same sex form strong peer bonds with each other following co-housing. Peer bonded rats exhibit affiliative displays toward their cagemates who are distressed whereas they exhibit agonistic behaviors toward strangers in these situations. Using innovative, genetic strategies in rats, we show that both oxytocin receptor (OXTR) bearing neurons and *Oxtr* signaling in the dorsal hippocampus are essential for peer bonds to form. Together, we have developed a new platform for studying peer bonding and demonstrate a neural pathway that governs this behavior.

## Introduction

Social attachments are a defining feature of social animals (Feldman, 2017; Lim and Young, 2006; Seyfarth and Cheney, 2012). Attachments among parents and progeny (parent-infant bonding), sexual partners (pair bonding), and non-related adults who are not sexually engaged (peer bonding) are the most common forms of social relationships (Feldman, 2017). While the neural mechanisms of the first two bonding types have been well studied (Carter et al., 1997; Feldman, 2007; Gobrogge and Wang, 2016; Goldberg, 1983; Johnson and Young, 2015), few studies have been done to understand the neural and molecular mechanisms of peer bonding due to the lack of appropriate experimental models (Schweinfurth et al., 2017). Nevertheless, both humans and other social animals benefit from close and enduring peer social relationships, as these relationships may help increase reproduction and survivability, and reduce stress and disease risk (Berkman et al., 2004; Cameron et al., 2009; Hill et al., 2009; Holt-Lunstad et al., 2010; Kohn, 2017; Silk, 2007). In addition, deficits in developing or maintaining friendships are common in psychiatric patients with disorders such as autism spectrum disorder (ASD) (Bauminger et al., 2008; Jost and Grossberg, 1996; Porcelli et al., 2019; Tough et al., 2017; Umberson and Karas Montez, 2010), highlighting an urgent need for an animal model to study the neural and molecular mechanisms of peer bonding.

Previously, social novelty assay was commonly used to evaluate social recognition ability which is a prerequisite of peer bonding (Raam et al., 2017). However, it’s essential to note that social recognition only reflects the episodic memory, offering a partial perspective on the multidimensional nature of peer social behaviors. Therefore, there is a pressing need for the development of an experimental model that can comprehensively reveal the features of peer social behaviors. This model should have the capability to effectively delineate the effect of the time accumulation on peer bonding under different contexts. The level of intimacy, which could be reflected by the intensity of social interactions such as affective touch, indicates the strength of peer bonding (Dunbar, 2010; Lackey and Williams, 1995; Nummenmaa et al., 2016). Affiliative social displays, like allogrooming, serve as comforting prosocial behaviors that can alleviate stress in the recipient (Hart and Hart, 1992; Jablonski, 2021; Liu et al., 2022; Rault, 2019; Spruijt et al., 1992; Wu et al., 2021), and have essential roles in building and strengthening social bonds throughout a wide range of species (Cameron et al., 2009; Dunbar, 2010; Silk et al., 2009; Val-Laillet et al., 2009). Indeed, an elevated allogrooming behavior toward distressed partners suggests peer caring in rodents (Burkett et al., 2016; Stieger et al., 2017; Zhang et al., 2022). Conversely, aversive social touch often triggers aggression and occurs in the context of social interactions with unfamiliar individuals (Saarinen et al., 2021). Thus, peer caring extent might be a breakthrough point to explore the features of peer bonding.

When opting for an appropriate animal model, it is imperative to consider the selection of rodent species due to the distinctions in social behavior between laboratory mouse and rat strains (Netser et al., 2020). Rats are widely used in neuroscience research with pharmacological and viral manipulation strategies (El-Ayache and Galligan, 2020). Living in large colonies, rats perform complex social behaviors to maintain their social structures within groups (Schweinfurth, 2020) and manifest prosocial behaviors including cooperation (Jiang et al., 2021; Schweinfurth and Taborsky, 2018) and helping (Bartal et al., 2011; Mason, 2021; Sato et al., 2015). Considering that social interactions between male mice are usually antagonistic (Weber et al., 2022), rats are more suitable to study peer bonding.

The oxytocin system is well-established for its role in promoting social interactions, including parent-infant interactions and pair bonding (Benjamin and Neumann, 2018; Insel, 2010; Marlin et al., 2015; Valery and Ron, 2018; Walum and Young, 2018). In the social interactions among same-sex peer mice, social recognition is also reported to be regulated by the hippocampal oxytocin receptor (OXTR) (Lin et al., 2018; Lin and Hsu, 2018; Pan et al., 2022; Raam et al., 2017; Tsai et al., 2022). Interestingly, a recent research has unearthed a disparity in the necessity of *Oxtr* function for pair bonding and parental behaviors in prairie voles (Berendzen et al., 2023), implying potential species-specific variations in the roles played by the oxytocin system in social bonding. Given that the precise mechanisms governing *Oxtr*’s involvement in animal bonding processes remain largely unknown, and we are still at an early stage of assessing oxytocin-based therapy for psychiatric disorders (e.g., autism) due to the mixed clinical trial results (Young and Barrett, 2015), it is important to ascertain whether and how hippocampal *Oxtr* plays a role in mediating peer bonding.

Here, we developed a stress-induced behavioral paradigm to investigate the neural basis of peer bonding establishment with adult peer (same-age, -sex) SD rats. We found that rats formed a close familiarity-based relationship that enabled care-taking for each other when facing distress. By contrast, lack of familiarity led to agonistic interactions between individuals under a similar condition. Based on the affiliative and agonistic actions induced by either forced swimming (FSW) or pain, we further demonstrated that two weeks was an adequate duration for rats to form peer bonding. With viral manipulation and knocking-out *Oxtr* in dorsal hippocampus, we found that dorsal hippocampal *Oxtr* was required for new peer bonding establishment but not maintenance.

## Results

### Rats exhibit close social relationships with their familiar peers in distress

Previous studies have demonstrated that stressors including pain, foot shock, forced swimming (FSW), and acute restraint, increase peer-caring behaviors such as allogrooming in rodents (Burkett et al., 2016; Du et al., 2020; Li et al., 2018; Lu et al., 2018; Wu et al., 2021). To explore whether adult rats can form peer bonding, we established a novel paradigm with FSW-induced social interaction on demonstrator-observer pairs with different levels of familiarity (Fig. 1A and Methods). In line with existing studies suggesting the formation of strong social bonds among familiar female individuals (Cameron et al., 2009; Christov-Moore et al., 2014), we initially recorded agonistic and affiliative behaviors during the female-female social interaction section (Fig. 1B and Movie S1). We found that behaviors observed between stranger rats differed from those between co-housed rats (Fig. 1C and Movie S2). Stranger rats exhibited more defensive behaviors and a higher aggressive ratio than rats co-housed for more than one week (Fig. 1D-1E), while affiliative behaviors were infrequent in stranger rats (Fig. 1F). The social behavior correlations also demonstrated differences between interactions in stranger rats and cagemate rats (Fig. S1A-S1B). Close relationships form between strangers after a certain amount of time (Levinger, 1980). As affiliative behaviors like allogrooming and crawling on top increased significantly after rats were kept together for two weeks, we reasoned that two weeks is sufficient for adult female rats to form close peer bonding.

**Fig. 1.**
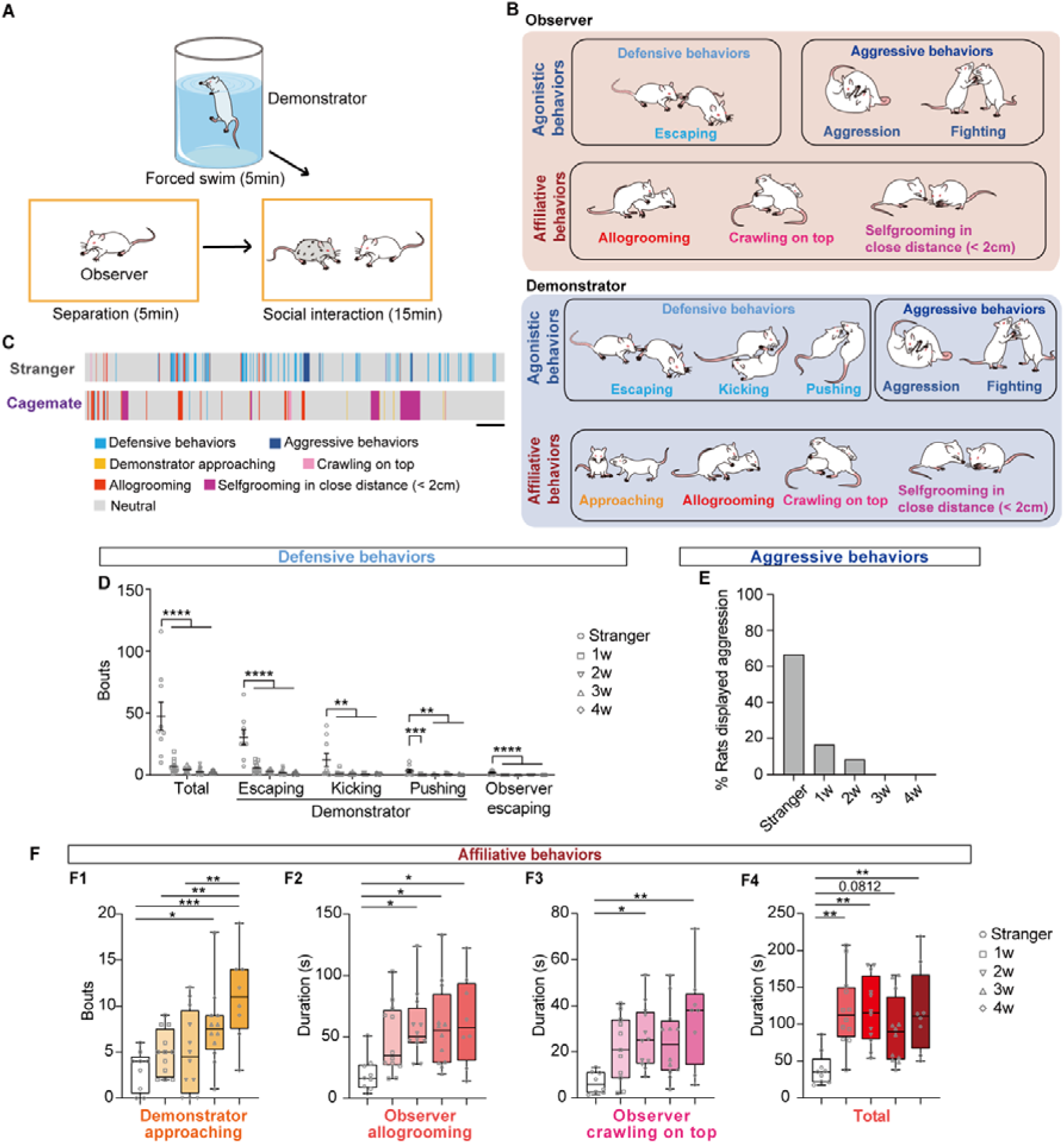
Rats exhibit close social relationships with their familiar peers in distress. (**A**) Schematic of forced-swimming consolation experiment setup. (**B**) Agonistic and affiliative behavior types during the experiment. (**C**) Behavioral description in two example sessions. Colors denote the different behaviors, as labeled in the key. Scale bar, 1 min. (**D**) Comparison of defensive behaviors in female rats co-housed for different durations. n_stranger_ = 9, n_1w_ = 12, n_2w_ = 12, n_3w_ = 12, n_4w_ = 8, One-way analysis of variance (ANOVA) followed by Tukey’s multiple comparisons test, Mean ± SEM, \*\**P* < 0.01, \*\*\**P* < 0.001, \*\*\*\**P* < 0.0001. (**E**) Ratio of aggressive behaviors in each group. n_stranger_ = 9, n_1w_ = 12, n_2w_ = 12, n_3w_ = 12, n_4w_ = 8. (**F**) Comparison of affiliative behaviors in female rats co-housed for different durations. n_stranger_ = 9, n_1w_ = 12, n_2w_ = 12, n_3w_ = 12, n_4w_ = 8, One-way ANOVA followed by Tukey’s multiple comparisons test, \**P* < 0.05, \*\**P* < 0.01, \*\*\**P* < 0.001.

The increased corticosteroid (CORT) level indicated elevated stress induced by a 5-min FSW (Fig. S1C). Additionally, observer CORT levels increased significantly after social interaction with its distressed cagemate (Fig. S1D), indicating a transfer of stress between peer-bonded rats. Interestingly, elevated CORT levels were also found in observers after interacting with distressed strangers. However, this increase was unlikely due to emotional contagion, as stranger observer CORT levels were also high after interactions with control strangers (Fig. S1D).

The social behaviors between male rats were also examined (Fig. S2). Similarly, stranger demonstrators displayed significantly more defensive behaviors (Fig. S2B), while affiliative behaviors between stranger male rats were remarkably lower than those between male rats co-housed for two weeks (Fig. S2D-S2H), further demonstrating that two weeks is adequate for rats of both sexes to form adult peer bonding.

### Pain induced similar features of adult peer bonding as FSW-induced distress

To determine whether peer bonding modulates social behaviors in different contexts, female rats were subjected to acute pain induced by injecting 2.5% formalin into the right hind paws of demonstrators (Fig. 2A); this acute pain causes physiological suffering, whereas FSW-induced stress is an affective state (Du et al., 2020). Consistent with previous results, stranger rats showed more defensive behaviors and higher aggressive ratio, while significantly less affiliative behaviors than rats co-housed for 2 weeks (Fig. 2B-2H). To examine whether pain can be transferred between individuals, a pain sensitivity test was performed with observers after a 15-min social interaction with painful stranger rats or cagemates (Fig. 2I). Cagemate observers exhibited a substantially lower paw withdrawal mechanical threshold (PWMT), whereas the PWMT of stranger observers was nearly 100% (Fig. 2J). This result suggests that pain can be transferred to cagemates but not strangers. Thus, both pain- and FSW-induced social interaction patterns demonstrate that adult SD rats can form a familiarity-based relationship that enables care-taking for each other when facing distress or in pain, and that two weeks of co-housing is sufficient for forming close peer bonding.

**Fig. 2.**
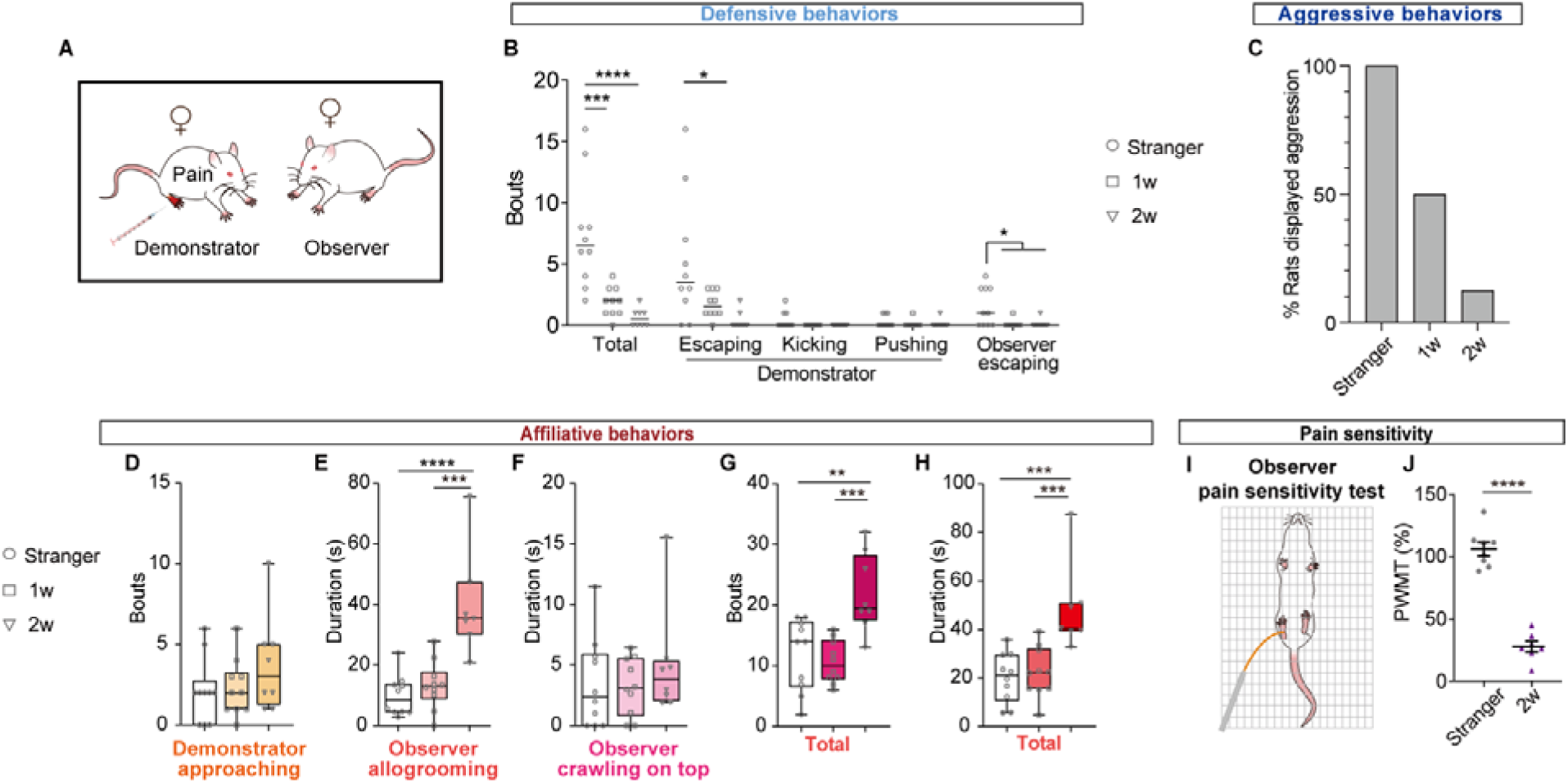
Rats exhibit close social relationships with their familiar peers in pain. (**A**) Experimental setup of an acute pain-induced stress model of female rats co-housed at different durations; resulting social behaviors were recorded. (**B**) Comparison of defensive behaviors in female rats co-housed for different durations. n_stranger_ = 10, n_1w_ = 10, n_2w_ = 8, One-way ANOVA followed by Tukey’s multiple comparisons test, \**P* < 0.05, \*\*\**P* < 0.001, \*\*\*\**P* < 0.0001. (**C**) Ratio of aggressive behaviors in each group. (**D-H**) Comparison of affiliative behaviors in female rats co-housed for different durations. n_stranger_ = 10, n_1w_ = 10, n_2w_ = 8, One-way ANOVA followed by Tukey’s multiple comparisons test, \*\**P* < 0.01, \*\*\**P* < 0.001, \*\*\*\**P* < 0.0001. (**I**) Schematic diagram of observer pain sensitivity test. (**J**) Comparison of paw withdrawal mechanical thresholds (PWMT) from stranger and familiar rats. PWMT (%) = PWMT_post-treatment_ / PWMT_baseline_ × 100. n_stranger_ = 8, n_2w_ = 7, unpaired Student’s *t*-test, Mean ± SEM, \*\*\*\**P* < 0.0001.

### Dorsal hippocampal neural activities of observers are distinct when exploring distressed stranger or cagemate

Next, to evaluate the brain regions that might be involved in peer bonding formation, we analyzed the expression of FOS protein (the protein that is expressed by the *c-fos* gene) in female observer rats that interacted with the distressed cagemate or stranger. CA1, DG, and CA3_proximal_ had higher ratios of FOS^+^ neurons than that of CA2/CA3_distal_ when comparing the cagemate and stranger groups (Fig. 3A-3B), suggesting that sub-regions of dorsal hippocampus may have different roles in peer bonding regulation in rats.

**Fig. 3.**
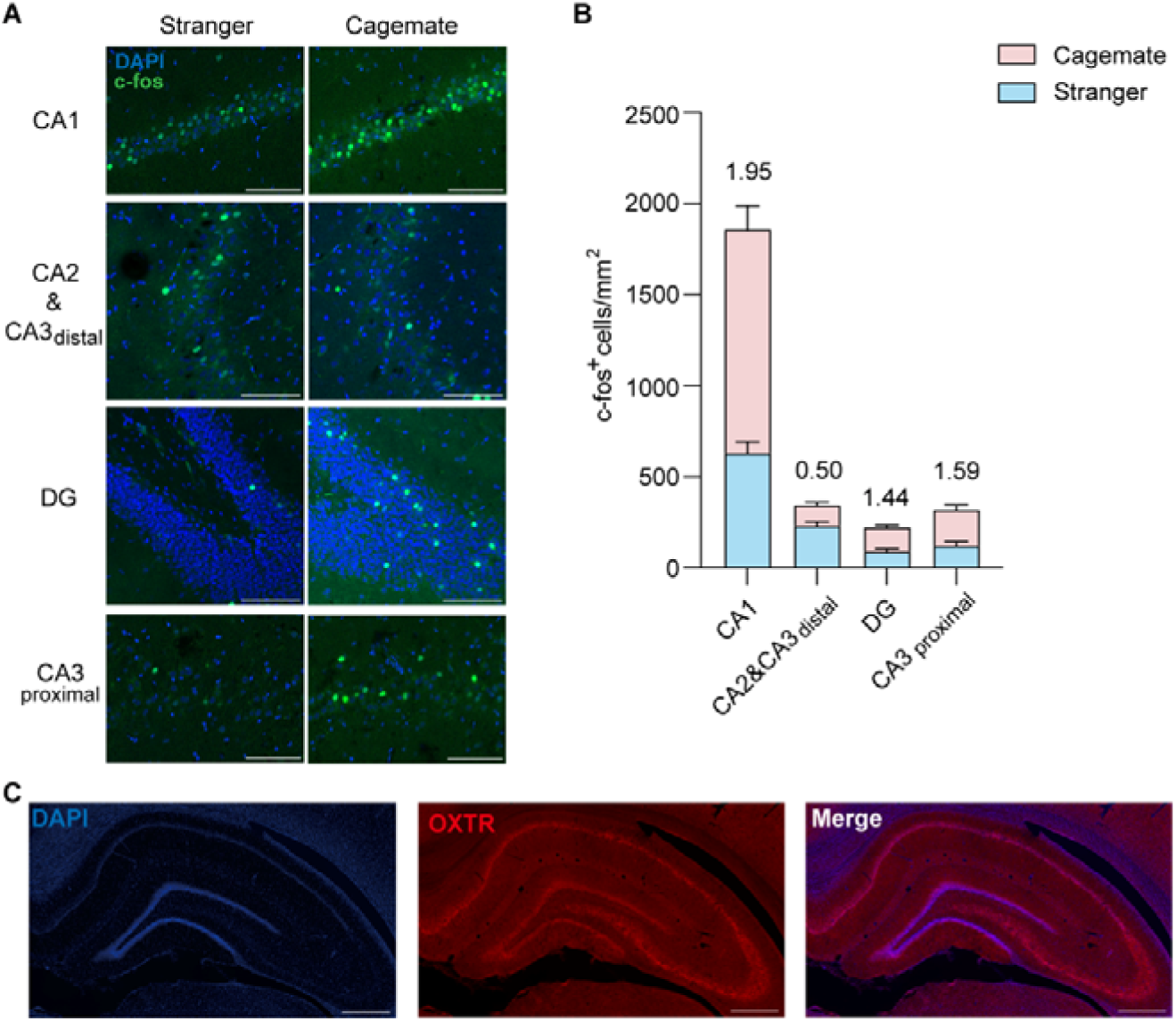
Dorsal hippocampal neural activities of observers are distinct when exploring distressed stranger or cagemate. (**A**) Representative images showing FOS^+^ neurons in the subregions of hippocampus from the cagemate and stranger observers (scale bars, 100 μm). (**B**) Histogram showing the number of FOS^+^ neurons per mm^2^ in CA1, CA2/CA3_distal_, DG and CA3_proximal_. The ratio of Cagemate/Stranger in each region is indicated on the up side of the bar chart. N_total_=18 (3 rats and 6 sections for each rat). (**C**) Representative images of OXTR expressions in dorsal hippocampus. Scale bar, 500 μm.

### Ablating dorsal hippocampal OXTR^+^ neurons impede peer bonding establishment

The oxytocin system was reported to play a well-established role in promoting social interactions, including parent-infant interactions and pair bonding (Benjamin and Neumann, 2018; Insel, 2010; Marlin et al., 2015; Valery and Ron, 2018; Walum and Young, 2018). Therefore, we investigated whether OXTR is also involved in the regulation of peer bonding. As OXTR was found to be expressed in dorsal hippocampus (Fig. 3C), we initially evaluated the role of dorsal hippocampal OXTR-expressing neurons in peer bonding regulation. We developed an ingenious experimental approach (Fig. S3A) to investigate the influence of these neurons on female rat peer bonding. Initially, OXTR-Cre adeno-associated virus (AAV) was bilaterally injected into the dorsal hippocampus of the target rat. One week later, the target rat was co-housed with a stranger (stranger A) for 2 weeks. Subsequently, the target rat received AAV carrying Cre-dependent taCasp3 (a genetically engineered caspase-3) (Yang et al., 2013) or EGFP into the dorsal hippocampus. After 1 week, a second phase of peer bonding was initiated by replacing stranger A with a new stranger rat (stranger B). Social behaviors with the previously co-housed distressed stranger A (old cagemate) and stranger B (new cagemate) were recorded respectively after these two distinct peer bonding phases (Fig. 4A and S3A, Methods). One week or five weeks after virus injection with taCasp3, EGFP^+^ OXTR^+^ neurons in the dorsal hippocampus (mainly in CA1, DG, and CA3_proximal_) were ablated (Fig. 4B and S3B-S3D), confirming the efficacy of taCasp3. Control rats exhibited similar defensive, aggressive and affiliative behaviors with old cagemates and new cagemates (Fig. 4C-4I). In contrast, rats with ablated dorsal hippocampal OXTR^+^ neurons displayed significantly lower affiliative behaviors with new cagemates compared to their old cagemates (Fig. 4E-4I). Notably, OXTR::taCasp3 rats exhibited normal affiliative behaviors toward the distressed old cagemates, indicating that dorsal hippocampal OXTR^+^ neurons ablation did not impair abilities to exhibit affiliative behaviors but directly impeded peer bonding establishment. The results of the elevated plus maze test (EPM) showed that anxiety levels were not influenced by ablating these neurons (Fig. S3E). To exclude the influence of social experience, we also compared social behaviors with SD rats that interacted with the old cagemate first or those that interacted with the new cagemate first. The results demonstrated that social behaviors were minimally affected by the social experience (Fig. S3F-S3N).

**Fig. 4.**
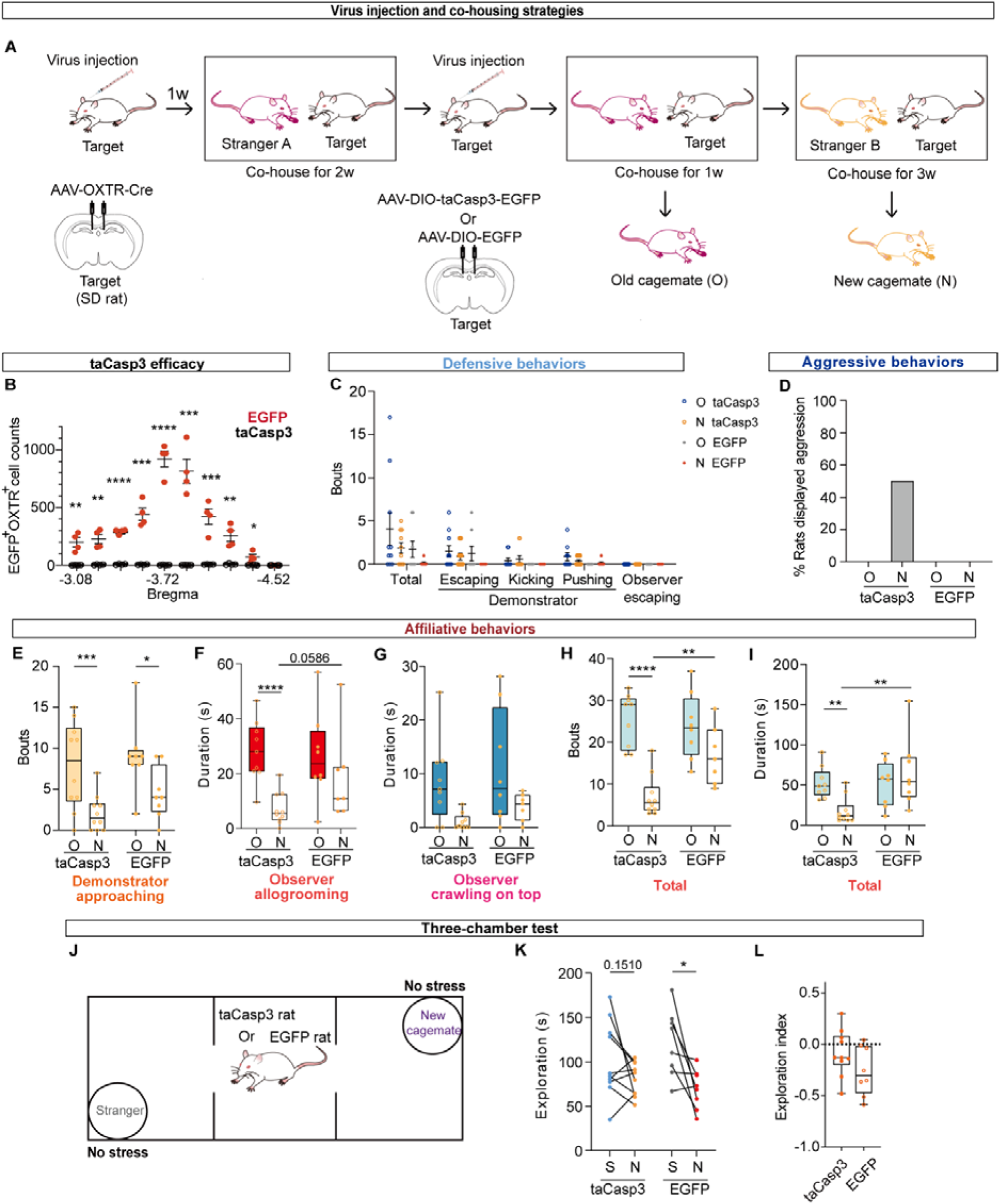
Ablating dorsal hippocampal OXTR^+^ neurons impede peer bonding establishment. (**A**) Virus injection and co-housing strategies. SD female rats (Target rats) were firstly injected with AAVs virus to express Cre in dorsal hippocampal OXTR^+^ neurons. After 1-week recovery and 2-week co-housing with SD female rats (Stranger A), they were randomly chosen as control target (EGFP) or treatment target (taCasp3) by injecting the corresponding Cre-dependent virus. After one more week co-housed with treatment or control rats, Stranger A rats were replaced with Stranger B rats for another 3 weeks. Stranger A and Stranger B were old cagemate (O) and new cagemate (N) of the target, respectively. (**B**) The efficacy of taCasp3 on ablating dorsal hippocampal OXTR-expressing neurons from Bregma -3.08 to Bregma -4.52 after virus were injected for 5 weeks. n=4 for each group, unpaired Student’s *t*-test, Mean ± SEM, \**P* < 0.05, \*\**P* < 0.01, \*\*\**P* < 0.001, \*\*\*\**P* < 0.0001. (**C**) Comparison of defensive behaviors of female rats in treatment (taCasp3) and control (EGFP) groups. n_taCasp3_=10 pairs for both old and new cagemate social interactions, n_EGFP_= 8 pairs for old cagemate social interaction and n_EGFP_= 7 pairs for new cagemate social interaction (an outlier was excluded), Mean ± SEM, Mixed-effects analysis, ns, no significance. (**D**) Ratio of aggressive behaviors in each group. n_taCasp3_=10 pairs, n_EGFP_= 8 pairs for both old and new cagemate social interactions. (**E-I**) Comparison of affiliative behaviors in female rats in treatment and control groups. n_taCasp3_=9-10 pairs, n_EGFP_= 7-8 pairs for old and new cagemate social interactions, Mixed-effects analysis, \**P* < 0.05, \*\**P* < 0.01, \*\*\**P* < 0.001, \*\*\*\**P* < 0.0001. (**J-L**) Social preference comparison of treatment and control rats. Experimental setup (**J**); statistical analysis of exploration time (**K**) and index (**L**), Exploration index = (Time_new cagemate_ − Time_stanger_)/(Time_new cagemate_ + Time_stanger_). n= 8–10, Paired Student’s *t*-test (**K**), unpaired Student’s *t*-test (**L**), \**P* < 0.05.

### Dorsal hippocampal *Oxtr* regulates peer bonding establishment

Considering that the slightly impaired social recognition ability induced by ablating dorsal hippocampal OXTR^+^ neurons (Fig. 4J-4L) could not fully elucidate the role of these neurons in peer bonding formation, we explored the role of dorsal hippocampal *Oxtr* in peer bonding by deleting *Oxtr* with CRISPR-Cas9 method. Initially, we generated a transgenic rat (LSL-Cas9) to facilitate the conditional expression of Cas9 in a Cre-dependent manner (Fig. 5A-5B and S4A). The decreased immunoreactivity (fluorescent intensity: control, 43.42 ± 0.5847; cKO, 28.50 ± 3.596; unpaired Student’s *t*-test, p=0.0149, n=3 control and 3 cKO) of *Oxtr* in female dorsal hippocampus validated the efficacy of the conditional knockout strategy (Fig. 5C-5D). Consequently, the *Oxtr* cKO female rats displayed significantly reduced affiliative behaviors toward the distressed new cagemates compared to their interactions with the old ones (Fig. 5E-5K). Notably, the rats exhibited normal anxiety levels (Fig. S4B) and social recognition ability (Fig. 5L-5N). These findings provide further evidence highlighting the essential role of *Oxtr* in dorsal hippocampus for peer bonding establishment, but not maintenance.

**Fig. 5.**
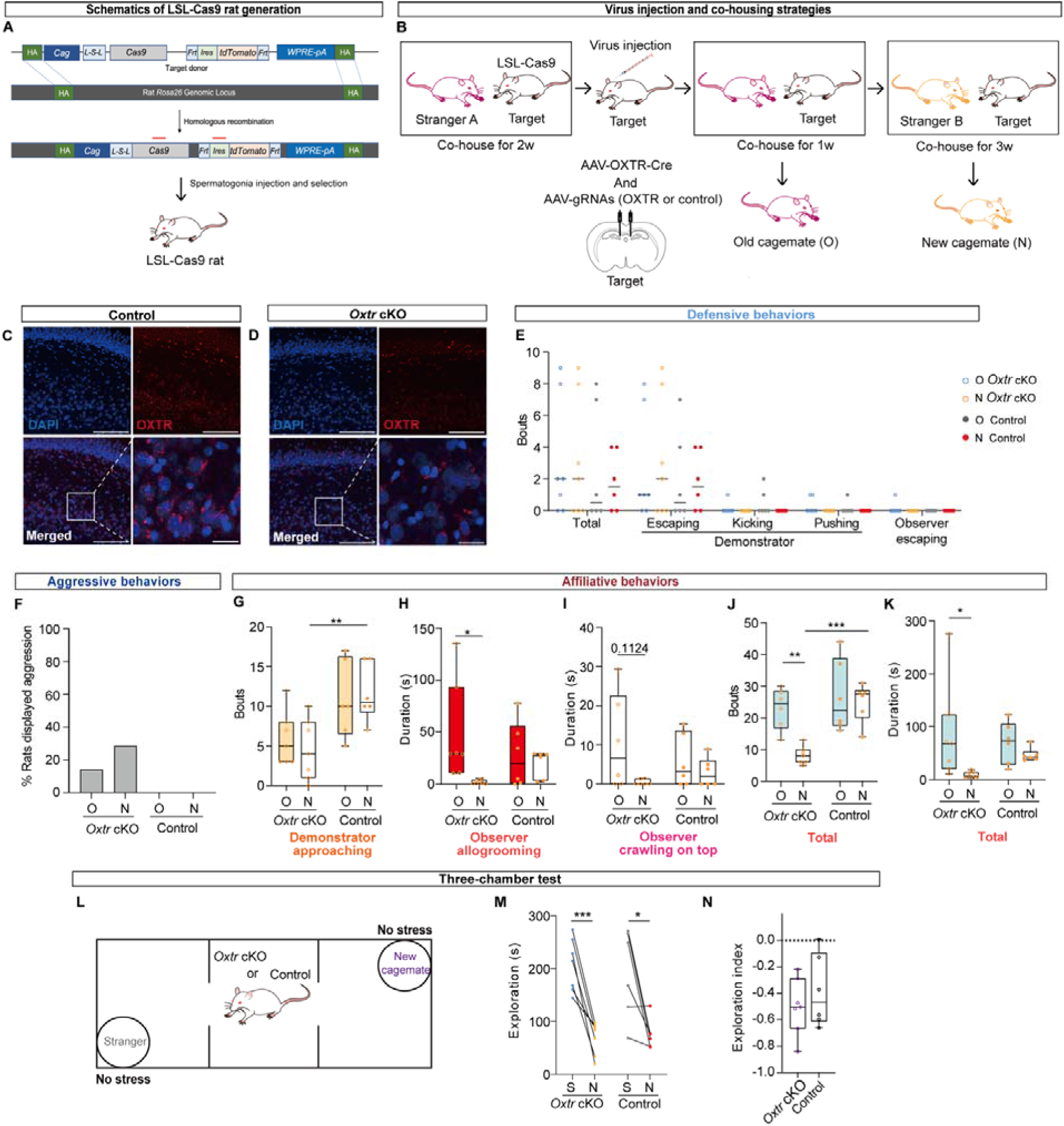
Dorsal hippocampal *Oxtr* regulates peer bonding establishment. (**A**) Schematics of generating LSL-Cas9 rat. Red bars indicate the locations of primers for genotyping. (**B**) Virus injection and co-housing strategies. (**C-D**) Representative images of OXTR expressions in dorsal hippocampus (DG and CA3_proximal_) of control rats (**C**) and *Oxtr* conditional knockout rats (**D**) after viruses were injected for 5 weeks. Scale bar, 100 μm (scale bar: 20 μm for the enlarged images). (**E**) Comparison of defensive behaviors of female rats in treatment (*Oxtr* cKO) and control groups. n_oxtr cKO_=7 pairs, n_control_=6 pairs, Two-way repeated ANOVA followed by Šídák’s multiple comparisons test, no significance. (**F**) Ratio of aggressive behaviors in each group. n_oxtr cKO_=7 pairs, n_control_=6 pairs for both old and new cagemate social interactions. (**G-K**) Comparison of affiliative behaviors in female rats in treatment and control groups. n_oxtr cKO_=6-7 pairs, n_control_=6 pairs for old and new cagemate social interactions, Two-way repeated ANOVA followed by Šídák’s multiple comparisons test (**G-H**, **J-K**), Mixed-effects analysis (**I**), \**P* < 0.05, \*\**P* < 0.01, \*\*\**P* < 0.001. **L-N**, Social preference comparison of treatment and control rats. Experimental setup (**L**); statistical analysis of exploration time (**M**) and index (**N**), Exploration index = (Time_new cagemate_ − Time_stanger_)/(Time_new cagemate_ + Time_stanger_). n= 6–7, Paired student’s *t*-test (**M**), unpaired Student’s *t*-test (**N**), \**P* < 0.05, \*\*\**P* < 0.001.

## Discussion

Here, our study addresses a fundamental gap in our understanding of the neural and molecular basis governing peer bonding in adult rodents. We have developed a stress-induced experimental model to study adult peer bonding and demonstrated that a two-week duration was adequate for rats to form this relationship. Furthermore, we established that dorsal hippocampal *Oxtr* is essential for new peer bonding establishment but not maintenance.

Our study satisfied the characteristics of the conceptual framework on social bonding developed by Lim and Young (Lim and Young, 2006). First, the individual must be motivated to approach and engage with another conspecific. In the context of rodent rats, it is well-established that they exhibit a strong inclination to engage in social interactions with unfamiliar conspecifics. Second, the animals must be able to identify the individual based on social cues through the formation of social memories. In our experiments, we employed the three-chamber test to demonstrate the capacity of rats to differentiate between conspecifics with varying degrees of familiarity. Finally, a bond can form when given appropriate conditions, leading to preferential interaction with that individual. In our investigation, we identified two crucial initial conditions for the formation of peer bonding: a 2-week duration of co-housing and the intact function of dorsal hippocampal *Oxtr*. Additionally, our results indicate that the distressed stranger always repelled the observer while the distressed cagemate accepted more consolation behaviors from its partner. Thus, our study satisfied the three characteristics of the conceptual framework on social bonding developed by Lim and Young.

Previously, affiliative behaviors, especially allogrooming behavior, have primarily been used as bonding indicators. Here, we not only expanded affiliative behavior types, but also added agonistic acts (including defensive and aggressive behaviors) of both demonstrator and observer rats, enabling a more comprehensive measurement of peer bonding. These multidimensional social behaviors increase the credibility of our investigation into social bonding dynamics. Furthermore, both the affiliative and agonistic social displays showed consistent patterns when the peer-bonded rat is in distress following forced swimming or is in pain following formalin injection into a paw. In other words, the experimental rat can generalize its affiliative or agonistic behaviors beyond one context.

In the realm of rodent prosocial interaction research, a longstanding question has been never asked and unexamined: the rationale behind the prerequisite that rodents employed in prosocial investigations should cohabitate as cagemates for several weeks before behavioral assessments. Our findings, demonstrating that a two-week cohabitation period is adequate for rats to establish a relatively stable peer bonding relationship, now provide a definitive answer to this hitherto unaddressed inquiry. While our results have underscored the requisite duration for rats to form such stable peer bonds, further investigations are warranted to pinpoint the precise temporal dynamics of this bonding event. Additionally, exploring whether these dynamics exhibit sexual dimorphism is essential, as behavioral nuances differ between male and female rats.

It is controversial that the oxytocin system is involved in mediating pair bonding and parental care, since prairies voles with null *Oxtr* were reported to show normal social attachment recently (Berendzen et al., 2023). The observed disparities in phenotypes could potentially be attributed to species-specific differences in the oxytocin system’s functionality. Differences in the localization of OXTR expression were observed, with mice displaying hilus-based expression in DG (Raam et al., 2017) and rats showing expression in the granular cell layer (Fig. 3C). Furthermore, previous study in mice demonstrated that DG hilar *Oxtr* is necessary for social discrimination (Raam et al., 2017), while we found that rats with *Oxtr* specially knocked out in dorsal hippocampus (mainly in CA1, DG, and CA3_proximal_) can still distinguish strangers from their new cagemates, but their ability to form new peer bonding was impaired. When considering the disparities in both OXTR expression and function evident across distinct rodent species, it becomes plausible to entertain the idea that the rapid evolution of these neural pathways has contributed to the divergence in neural mechanisms governing social interaction among rodents.

It has long been believed that OXTR signaling in CA2/CA3 of hippocampus is crucial for regulating social recognition in same-sex peer mice (Lin et al., 2018; Lin and Hsu, 2018; Pan et al., 2022; Raam et al., 2017; Tsai et al., 2022). Here, we found that, when comparing the FOS^+^ neuron expression in cagemate and stranger groups, the ratio of CA2/CA3_distal_ was lower than 1 (Fig. 3B), suggesting that neurons in this region may be activated when rats encountering a distressed stranger. Our results in rats align with previous findings in mice, indicating the potential preserved role of social recognition-related signaling in CA2/CA3 across species. Besides, the different ratio patterns of FOS^+^ neurons in subregions of rat dorsal hippocampus in our study shed light on the possibility that dorsal hippocampal OXTR may have different divisions of labor on peer bonding regulation in rats.

In our investigation, we observed that rats with a specialized knockout of *Oxtr* in dorsal hippocampus (mainly in CA1, DG, and CA3_proximal_) displayed impaired bonding with their distressed new cagemates, but their affiliative behaviors remained high with their distressed old cagemate rats. This outcome effectively rules out the possibility that dorsal hippocampal *Oxtr* regulates peer bonding directly without influencing the rats’ capacity for social behaviors. As we only measured the bonding level through the behavioral paradigm, future studies with techniques such as two-photon imaging could provide in-depth insights into the physiological characteristics of neurons in rats. Given the distinct behavioral responses observed when rats interacted with distressed strangers, old cagemates and new cagemates, it is conceivable that dorsal hippocampal neurons may exhibit varying response patterns in these contexts.

Oxytocin has long been recognized as a potential ASD treatment (Andari et al., 2010; Guastella et al., 2010; Ooi et al., 2017), and certain autistic patients have been reported to have genetic variants in the *Oxtr* gene (Campbell et al., 2011; Lerer et al., 2008; LoParo and Waldman, 2015; Wu et al., 2005). Moreover, individuals with ASD encounter challenges in establishing relationships and bonding with unfamiliar individuals. With the innovative rat model, we establish that dorsal hippocampal *Oxtr* is required for peer bonding formation, suggesting the potential avenue for treating autism patients with oxytocin system by leveraging the neural mechanisms underpinning peer bonding. While we found the necessity of *Oxtr* in peer bonding establishment, it remains unknown whether *Oxtr* alone is sufficient to enhance the bonding between peer strangers. As the application of oxytocin therapy for autism is fraught with complexities, there is a compelling need to delve deeper into the neural and molecular mechanisms that underlie the regulation of peer bonding.

## Methods

### Rat lines and animal care

Two rat strains were used in the experiments. Sprague Dawley (SD) rats, purchased from Beijing Vital River Laboratories Animal Center, were used for the exploration of peer bonding formation time. The Cre-dependent Cas9 rat line (LSL-Cas9) was generated using a CRISPR/Cas9-based approach. Rats were sex- and age-matched (within one week) and housed 2–4 per cage in standard laboratory conditions under a 12-h light/dark cycle with *ad libitum* access to food and drinking water except during experimental sessions. All experiments were conducted with adult rats (8–15 weeks) and rats used for behavioral tests were 12–15 weeks old. Behavioral testing occurred during the dark period. All experiments were performed in accordance with procedures approved by the Institutional Animal Care and Use Committee at the Institute of Basic Medical Sciences, Chinese Academy of Medical Sciences.

### Adeno-associated virus (AAV)

rAAV-ef1α-DIO-taCasp3-TEVp-P2A-EGFP-WPRE-hGH polyA (2.32×10^12^ particles/ml), rAAV2/9-ef1α-DIO-EGFP-WPRE-hGH polyA (2.56×10^12^ particles/ml), rAAV-OXTR-Cre-WPRE-pA (2×10^12^ particles/ml), rAAV-U6-sgRNA(Scramble)-U6-sgRNA(Scramble)-CMV-EGFP-WPRE-polyA (2.5×10^12^ particles/ml), and rAAV-U6-sgRNA1(*Oxtr*)-U6-sgRNA2(*Oxtr*)-CMV-EGFP-WPRE-polyA (2.5×10^12^ particles/ml) were purchased from BrainVTA (Wuhan, China).

### Generation of LSL-Cas9 transgenic rat

The LSL-Cas9 rat line was developed by Shanghai Model Organisms Center, Inc. This rat line was generated with CRISPR/Cas9 system by knocking in Cas9 cassette into the SD rat *Rosa26* locus, which is the most frequently used locus to produce ubiquitous or controlled expression of a gene of interest in rodents. Briefly, targeting vector containing the following components were constructed: CAG-LSL-Cas9-FRT-IRES-tdTomato-FRT-WPRE-polyA. Cas9 mRNA was *in vitro* transcribed with mMESSAGE mMACHINE T7 Ultra Kit (Ambion, TX, USA) according to the manufacturer’s instructions, and subsequently purified using the MEGAclear^TM^ Kit (ThermoFisher, USA). 5’-GGAGCCATGGCCGCGTCCGG-3’ and 5’-GGACGGGCGGTCGGTCTGAG-3’ were chosen as Cas9 targeted guide RNAs (sgRNAs) and in vitro transcribed using the MEGAshortscript Kit (ThermoFisher, USA) and subsequently purified using MEGAclear^TM^ Kit. The donor vector with sgRNA and Cas9 mRNA was microinjected into the fertilized eggs of SD rats. PCR genotyping (Forward: 5’-CAACTCACAACGTGGCACTG-3’, Reverse: CCTGTACGAGACACGGATCG) and sequencing confirmed appropriate targeting to the *Rosa26* locus.

### Stereotactic surgery

SD rats (aged 8 weeks) were anesthetized with isoflurane (3-4% for induction, 1-2% for maintenance) and mounted on a stereotaxic frame (RWD, 68027). Small holes were drilled bilaterally after the skull was exposed, and were injected with virus at a rate of 30 nl/min (150nl per site) by using a pulled, fine glass micropipette (World Precision Instruments, Cat#504949). The stereotaxic coordinate of the dorsal hippocampus used were as follows (measured from Bregma in mm): anteroposterior -3.72, mediolateral ±2.2, and dorsoventral -3.2 determined according to Paxinos and Watson’s *The Rat Brain in Stereotaxic Coordinates* atlas. After completion of the injection, the micropipette was left on the site for an additional 10 min to allow the diffusion of the virus and the electrode was withdrawn slowly before skin was sutured. After surgery, rats were returned to their home cage and monitored during the recovery.

### Social interaction after forced swimming

To examine social interactions between rats, we adapted a paradigm that was modified from a previous study in mice (Wu et al., 2021). The observer was isolated for 5 min, while the demonstrator was placed into a beaker containing room temperature (20–23 LJ) water. The demonstrator was then removed from the beaker and placed together with the observer for 15 min, during which time their social interactions were recorded with an infrared camera installed above the cage (40 cm × 25 cm × 20 cm).

The two main behavioral aspects were recorded and statistically analyzed: 1) agonistic acts, including defensive behavior (i.e., kicking, pushing, and escaping) and aggressive behavior (i.e., aggressive allogrooming and fighting); 2) affiliative behaviors, including allogrooming (observer or demonstrator licks the fur on the back or head of the other rat for more than 1 s), crawling on top (observer or demonstrator crawls onto the back of the other rat laterally), demonstrator approaching (demonstrator comes to the near side of the observer proactively), and selfgrooming in close distance (rats < 2 cm apart while selfgrooming). Other behaviors were categorized as neutral behaviors and include exploring and investigating the home cage, walking, and selfgrooming. Since some behaviors—such as defensive behaviors and demonstrator approaching—occur in a split second, counts were recorded for these behaviors instead of durations. The total affiliative behavior bouts include bouts of demonstrator approaching, observer and demonstrator allogrooming, demonstrator and observer crawling on top, and demonstrator and observer selfgrooming in close distance (< 2 cm); total affiliative behavior durations include durations of observer and demonstrator allogrooming, demonstrator and observer crawling on top, and demonstrator and observer selfgrooming in close distance (< 2 cm).

### Social interaction after formalin injection in hind paws

The pain consolation paradigm was modified from a previous study (Du et al., 2020). Briefly, rats were randomly chosen as observers or demonstrators. The demonstrator was muffled with a piece of towel for immobility and the right hind paw was injected with 100 μL 2.5% formalin via a microsyringe (Gaoge, China) before it was returned to the cage. Social behaviors, including agonistic acts and affiliative behaviors, were recorded for 15 min with an infrared camera pre-positioned at a top view over the cage.

### Quantitative pain sensory test with von Frey filaments

To evaluate the transfer of pain in stranger groups and familiar groups, the observer of each group was used for the mechanical pain sensitivity test based on previously described experimental procedures (Du et al., 2020; Yu et al., 2019). The equipment setup included a nontransparent plastic testing box (10 cm × 30 cm × 20 cm) placed on a supporting platform equipped with a metal mesh (pore size 0.5 cm × 0.5 cm). Von Frey Hairs (Cat. No. 37450-275, Ugo Basile, Italy) were used for the mechanical pain sensitivity test.

The observer was acclimated to the experimental environment for two days prior to the day of the experiment; this adaption process included handling, placing, and adapting to other objects in the behavioral testing room. Additionally, the observer received stimulus with an ascending series of calibrated von Frey filaments with intensities ranging from 39.2 mN to 588 mN in the plastic testing box during this two-day acclimation period. Each stimulation was repeated 10 times with an interval of at least 10 s, and the percentages of paw withdrawal were recorded. The mean of the smallest intensities that induced more than 50% paw withdrawal reflex during the two acclimation days was used to establish the baseline.

On the test day, the observer was placed into the plastic testing box after the 15-min social interaction with the distressed demonstrator. The percentages of post-treatment paw withdrawal were measured in response to increasing stimulation intensities from the von Frey filaments until the force was sufficient to elicit more than 50% paw withdrawal reflex. The data for paw withdrawal mechanical threshold (PWMT) was normalized using the following formula: PWMT(%) = PWMT_post-treatment_/PWMT_baseline_ × 100. All tests were performed in a blinded manner.

### Old cagemate and new cagemate experimental strategy

For experiments involving rats with ablated dorsal hippocampal OXTR-expressing neurons: To begin, we injected AAV-OXTR-Cre into dorsal hippocampus of SD rats (target rats) bilaterally to facilitate the expression of Cre in OXTR-expressing neurons. After the initial injection, each target rat was co-housed with a stranger rat (Stranger A) for a period of 2 weeks. Subsequently, the target rats received a second injection in the same dorsal hippocampal location, this time with AAV-DIO-taCasp3 or AAV-DIO-EGFP. Following the second injection, the target rats continued to cohabitate with Stranger A for an additional week. At this point, Stranger A was replaced with another stranger rat (Stranger B) which was co-housed with the target rat for another 3 weeks. Stranger A and Stranger B served as the old cagemate and new cagemate, respectively. The social behavioral assay was conducted on the following week. Old cagemates or new cagemates were designed as demonstrators, while the target rats served as observers.

For experiments involving rats with dorsal hippocampal *Oxtr* conditionally knocked-out: The experimental strategy was similar to the previous one. Briefly, LSL-Cas9 rats (target rats) were co-housed with Stranger A for 2 weeks initially. Following this, the target rats were injected with a combination of AAV-OXTR-Cre and AAV-Oxtr-gRNA (or AAV-control-gRNA) virus. The target rats continued to co-habit with Stranger A for an additional week after the viral injection. The subsequent procedures were consistent with the above-described strategies.

### Elevated plus maze test

The elevated plus maze (EPM) test was performed as previously described (Pellow et al., 1985). The plus-cross-shaped maze was custom-made of black Plexiglas consisting of two open arms (50cm × 10cm) and two enclosed arms (50cm × 10cm × 40cm) extending from a central square platform (10cm × 10cm) mounted on a wooden base raised 85cm above the floor. Rats were placed on the center square platform with their face toward an open arm and allowed to freely explore for 10min under a dimmed illumination (about 10 Lux). The behavior of the animals was videotaped by a camera (Shanghai XinRuan Information Technology Co., Ltd) and analyzed by Any-maze (Stoelting). The chamber was thoroughly cleaned with 70% ethanol, and then with paper towels moistened with distilled water. The apparatus would be dried with paper towels before each trial.

### Three-chamber test

The three-chamber test for discriminating cagemate from stranger rats was modified from a previous study (Nadler et al., 2004). The apparatus was a rectangular, three-chambered box fabricated from clear polycarbonate (120 cm × 40 cm × 30 cm) and equipped with dividing walls with retractable door-ways that allowed access into each chamber. Prior to the experiment, each observer was allowed to freely explore the three chambers twice for 10 min each time and a 2-h interval between each visit; meanwhile, each demonstrator was adapted to the wire cup (10 cm diameter, 30 cm height) twice for 10 min each time and a 2-h interval between each adaptation episode. On the day of the experiment, the observer rat was first placed into the center chamber with the other two chambers closed and allowed to freely explore for 10 min. The stranger and cagemate were placed in the wire cups prior to opening the doors of the two chambers. The observer was then allowed to explore all three chambers and its behavior was recorded for 10 min with a camera pre-positioned with a top view over the apparatus. The exploration index was calculated using the following formula: Exploration index = (Time_cagemate_ − Time_stanger_)/(Time_cagemate_ + Time_stanger_)

The chamber was thoroughly cleaned with 70% ethanol followed by paper towels moistened with distilled water. The apparatus was dried with paper towels before each trial.

### Serum CORT measurement

To measure the serum CORT levels, the rats were initially anesthetized with isoflurane immediately after experiencing the corresponding treatments, which included no treatment, 5-minute isolation, 5-minute forced swimming, and 15-minute social interaction with either distressed or non-distressed conspecifics. Subsequently, blood samples were collected from the tail vein without restraint within a 5-minute timeframe after the anesthetic induction. The rats used for the first blood collection were allowed to recover for at least two weeks until the second blood collection, ensuring that the stress from the blood collection did not influence subsequent behavioral parameters. Approximately 500 μL blood was collected from each rat and kept for 1 h at room temperature. The blood samples were centrifuged for 10 min at 3,000 rpm at 4 LJ, and serum was then collected from the stratified samples and stored at -80 LJ until use.

Before the CORT was measured, samples were treated with Bond Elut Plexa Solid Phase Extraction (SPE) cartridges (30 mg, 1 mL, Part No. 12109301, Agilent Technologies, Inc., CA, USA) and hormones were separated on an Infinity Lab Poroshell HPH-C8 column (2.1 × 50 mm, 2.7 μm, Agilent Technologies, Inc.). Liquid chromatography tandem mass spectrometric detection (LC-MS/MS) was performed on the 6495 Triple-Quadrupole LC/MS system (Agilent Technologies, Inc.) equipped with an ESI source. The LC1290 liquid chromatography system (Agilent Technologies, Inc.) was used to deliver the mobile phases with 0.1% formic acid (FA) in water (A) and methanol (B) at a flow rate of 0.3 mL/min. A sample volume of 50 μL was injected onto an HSS T3 LC column (1.8 mm, 200 AL, 2.1 mm I.D. × 100 mm, Waters, Belgium). Gradient LC flow started with 20% B, followed by a linear increase to 100% B in 4 min and held at 100% B for 2 min. The gradient was returned to 20% B in 0.01 min and held at 20% B for 2 min for column equilibration, for a total run time of 8 min. The sheath gas temperatures were set at 350 °C, and the sheath gas flow was set at 11 L/min. The mass spectrometer was operated in positive mode with a capillary voltage of 3,000 V. Multiple reaction monitoring (MRM) scan mode was used for the mass spectrometric detection and quantification of the corticosterone in positive mode. The dominant precursor ions were the molecular ion m/z 347.2 in ESI (+). The transition products measured in ESI (+) were m/z 329.2 (obtained by applying a collision energy (CE) of 12 V), and m/z 121 (CE 24V). MassHunter Workstation Software LC/MS Data Acquisition (Version B.07.01 Build 7.1.7112.0; Agilent Technologies, Inc.) was used for data acquisition, and Quantitative Analysis (Version B.07.00 Build 7.0.457.0; Agilent Technologies, Inc.) was used for data analysis.

### Immunohistochemistry

Rats were anesthetized with isoflurane and perfused with 0.01M PBS, followed by fixation of 4% paraformaldehyde (PFA). For FOS protein expression analyses, rats were sacrificed 90 min after completing the social interactions and their brains were quickly collected. Brains were collected and post-fixed in 4% PFA overnight at 4LJ, then were put into 20% and 30% sucrose at 4LJ for 2 days, respectively. Coronal sections were obtained at 16μm using a Leica CM3050 S cryostat. Floating sections were used for immunohistochemical labeling. Briefly, sections were washed in 1×PBS and blocked in 1×PBS containing 5% Albumin bovine V (BSA) and 0.5% Triton X-100 for 1.5-2h at room temperature. Then the sections were incubated in primary antibodies in blocking buffer containing 3% BSA and 0.3% Triton X-100 and shook for 2h at room temperature before stored at 4LJ overnight. Primary antibodies were used as follows: c-Fos (Rabbit, Synaptic Systems, Cat#226 308, 1:1000), OXTR (Rabbit, Alomone, Cat#AVR-013, 1:100) (Warfvinge et al., 2020). The following day, sections were washed thrice with 1×PBS and incubated with fluorescent-label-coupled secondary antibodies (Alexa 488 or 633-conjugated goat anti-rabbit IgG 1:1000; Invitrogen) for 2h at room temperature. After washing briefly, sections were stained with DAPI for 5min and mounted with mounting medium (Solarbio, Cat#S2100). Images were acquired using Zeiss LSM 980, Leica TCS SP8 gSTED or Nikon AXR laser scanning microscope at 20×.

### Quantification of FOS^+^ neurons and EGFP^+^OXTR^+^ neurons

Densities of c-Fos^+^ cells in hippocampal regions of SD rats were quantified using systematic optical density measurements by ImageJ, following a similar approach to a previously published protocol (Ghashghaei and Barbas, 2001). Brain slices, 40 μm in thickness, spanning from Bregma -2.76 to -3.96, were selected for analysis, with one brain slice chosen every 5 successive slices. The numbers of labeled neurons in hippocampal regions were estimated from counts of positively stained cells in 6 sections per brain region. Differences in cell counts were statistically evaluated using unpaired *t*-tests to examine the interactions among groups. Counting was performed by an investigator who was blinded to the treatment conditions.

For the assessment of EGFP^+^OXTR^+^ cell counts and fluorescence intensity, images were normalized to ensure uniform exposure time across all samples. Brain slices, 20 μm in thickness, spanning from Bregma -3.08 to -4.52, were selected for analysis, with one brain slice chosen every 8 slices. Utilizing Fiji (ImageJ), the mean fluorescent intensity (MFI) was measured within a designated region of interest (ROI, DG & CA3proximal) of 500×500 μm², along the same rostro-caudal axis for each image. The MFI of OXTR in each brain slice was then calculated using the formula: Final MFI = MFI of ROI − MFI of the background. Subsequently, the average Final MFI across brain slices within one rat was computed.

### Quantification and statistical analysis

All data were collected by experimenters who were blinded to the surgical treatments. The data in scatter dot plots are presented as mean ± SEM and data in box plots are displayed as the minimum (Min) to the maximum (Max) values. All statistical analyses were carried out using GraphPad Prism 9 (GraphPad Software, Inc.) and MATLAB (R2008b, MathWorks). Student’s *t* test, Wilcoxon test or Mann-Whitney test was used to evaluate the statistical significance between two datasets. For multiple comparisons, one-way analysis of variance (ANOVA), two-way ANOVA or Mixed-effects analysis was used. The statistical significance is indicated as follows: n.s., no significance; * p < 0.05, ** p < 0.01, *** p < 0.001, **** p < 0.0001. Detailed information for all statistical analyses including sample sizes, statistical test(s), and exact p values when p > 0.0001 are presented in Table S1. Example micrographs show representative results based on at least three independent biological samples.

## Acknowledgments

We thank Dr. Yi Rao for providing LSL-Cas9 rats. We thank Dr. Wei Zhang at Tsinghua university for valuable suggestions on the manuscript. We thank Dr. Chao Ma and Dr. Fan Liu at the Institute of Basic Medical Sciences Chinese Academy of Medical Sciences for help with pain sensitivity threshold test. We also thank Dr. Jianjun Zhang at Institute of Psychology, Chinese Academy of Sciences for providing Any-maze software. We thank National Center for Protein Sciences at Peking University in Beijing, China, for assistance with serum CORT measurement, acquiring confocal images, and making rat behavior graphs. This work was supported by grant 2021ZD0203300 from the Ministry of Science and Technology of the People’s Republic of China, grant 2021-I2M-1-034 from the Chinese Academy of Medical Sciences Innovation Fund for Medical Sciences, grant 3332021040 from Fundamental Research Funds for the Central Universities, grant 2060204 from State Key Laboratory Special Fund, grant KF-202302 from the Open Research Fund of the National Center for Protein Sciences at Peking University in Beijing and Y.H. is an awardee of Young Elite Scientist Sponsorship Program by Beijing Association for Science and Technology.

## Author information

Laixin Liu

Present address: School of Life Sciences, Peking-Tsinghua Center for Life Sciences, and Peking University McGovern Institute, Peking University, Beijing

Authors and affiliations

Department of Physiology, Institute of Basic Medical Sciences Chinese Academy of Medical Sciences, School of Basic Medicine Peking Union Medical College, Beijing

Yufei Hu, Wensi Li, Yinji Zhao, Yuying Liu, Wenyu Sun, Yi Yan, Laixin Liu & Pu Fan Chinese Institute for Brain Research, Beijing

Bowen Deng Contributions

Y.H., W.L., Y.Z., Y.L., Y.Y., and L.L. performed the experiments. B.D. contributed the generation of LSL-Cas9 rat. Y.H. and P.F. analyzed data and wrote the manuscript. Y.H. and W.S. made the figures. All authors discussed and commented on the manuscript.

Corresponding author

Correspondence to Pu Fan.

## Data and materials availability

All data is available in the main text and the supplementary materials.

