## Supplemental files for "Dorsal hippocampal oxytocin receptor regulates adult peer bonding in rats"

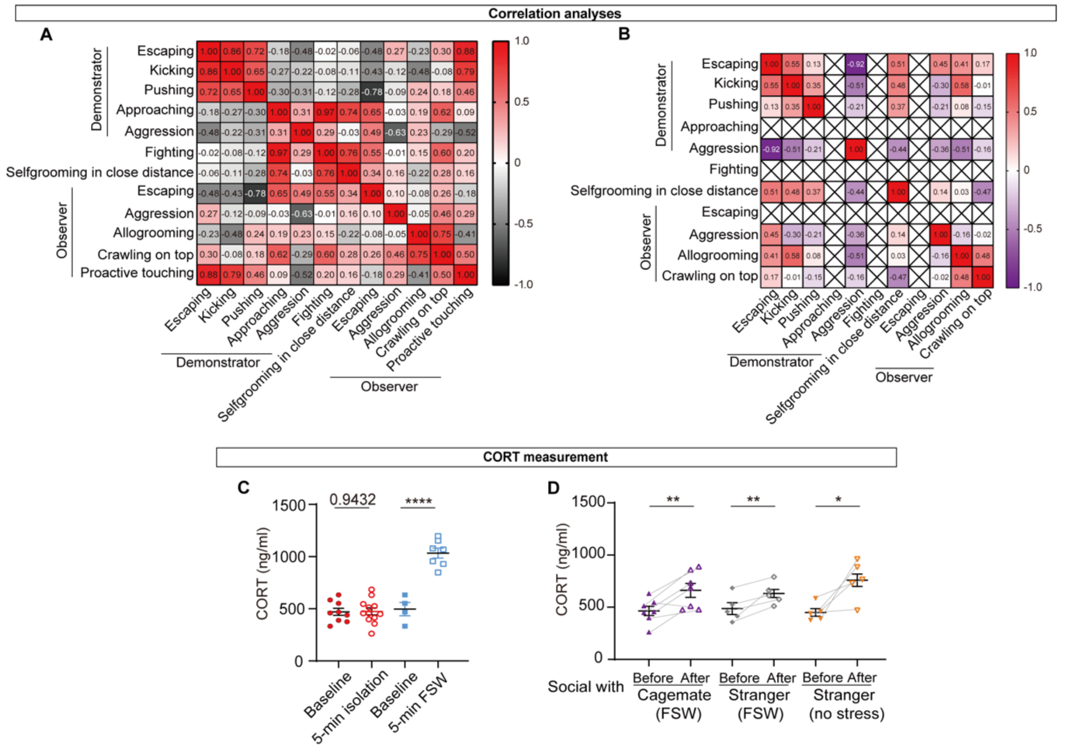


**Figure S1. Correlation analyses of social behaviors and Corticosteroid (CORT) levels of rats with different treatments.**

(**A**) Correlation analyses of social behaviors in stranger rats. Pearson's correlation, n = 9 pairs of female rats. (**B**) Correlation analyses of social behaviors in rats co-housed for 2 weeks. Pearson's correlation, n = 12 pairs of female rats. (**C**) Corticosteroid (CORT) levels of rats with different treatments. CORT levels from control rats (solid red) were not subjected to stress and served as the baseline, while CORT levels from rats isolated for 5 min were the treatment (open red circle). CORT levels of rats before (solid blue square, Baseline) and after a 5-min forced swimming (FSW) test (open blue square). (**D**) CORT levels of rats with different treatments and social interactions. Before (solid shapes) indicates rats after a 5-min isolation but prior to social interaction, and After (open shapes) indicates rats after a 15-min social interaction. Purple, CORT levels of rats before and after a 15-min social interaction with cagemates that suffered a 5-min FSW. Grey, CORT levels before and after a 15-min social interaction with strangers that suffered a 5-min FSW. Orange, CORT levels before and after a 15-min social interaction with strangers that had not been subjected to stress. n=4-12 (**C**), n = 5–7 (**D**), Unpaired Student's t-test (**C**), Paired Student's t-test (**D**), mean ± SEM, *P < 0.05, **P < 0.01, ****P < 0.0001.

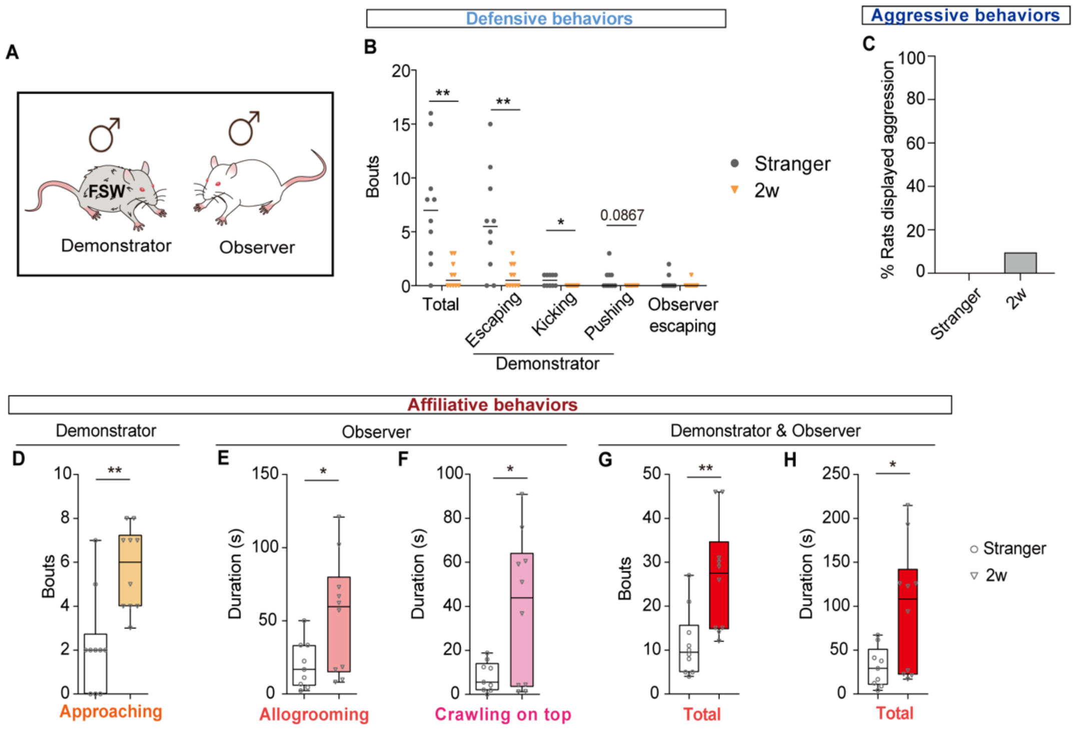


**Figure S2. Male rats exhibit close social relationships with their familiar peers in distress.** (**A**) Experimental setup. Social behaviors of stranger male rats and rats co-housed for 2 weeks (2w) were recorded. (**B**) Comparison of defensive behaviors in stranger male rats and rats co-housed for 2 weeks. nstranger = 10, n2w = 10, Mann-Whitney test, *P < 0.05, **P < 0.01. (**C**) Ratio of aggressive behaviors in each group. (**D-H**) Comparison of affiliative behaviors in stranger male rats and rats co-housed for 2 weeks. nstranger = 10, n2w = 10, Mann-Whitney test (**D**), Unpaired Student's t-test (**E-H**), *p < 0.05, **p < 0.01.

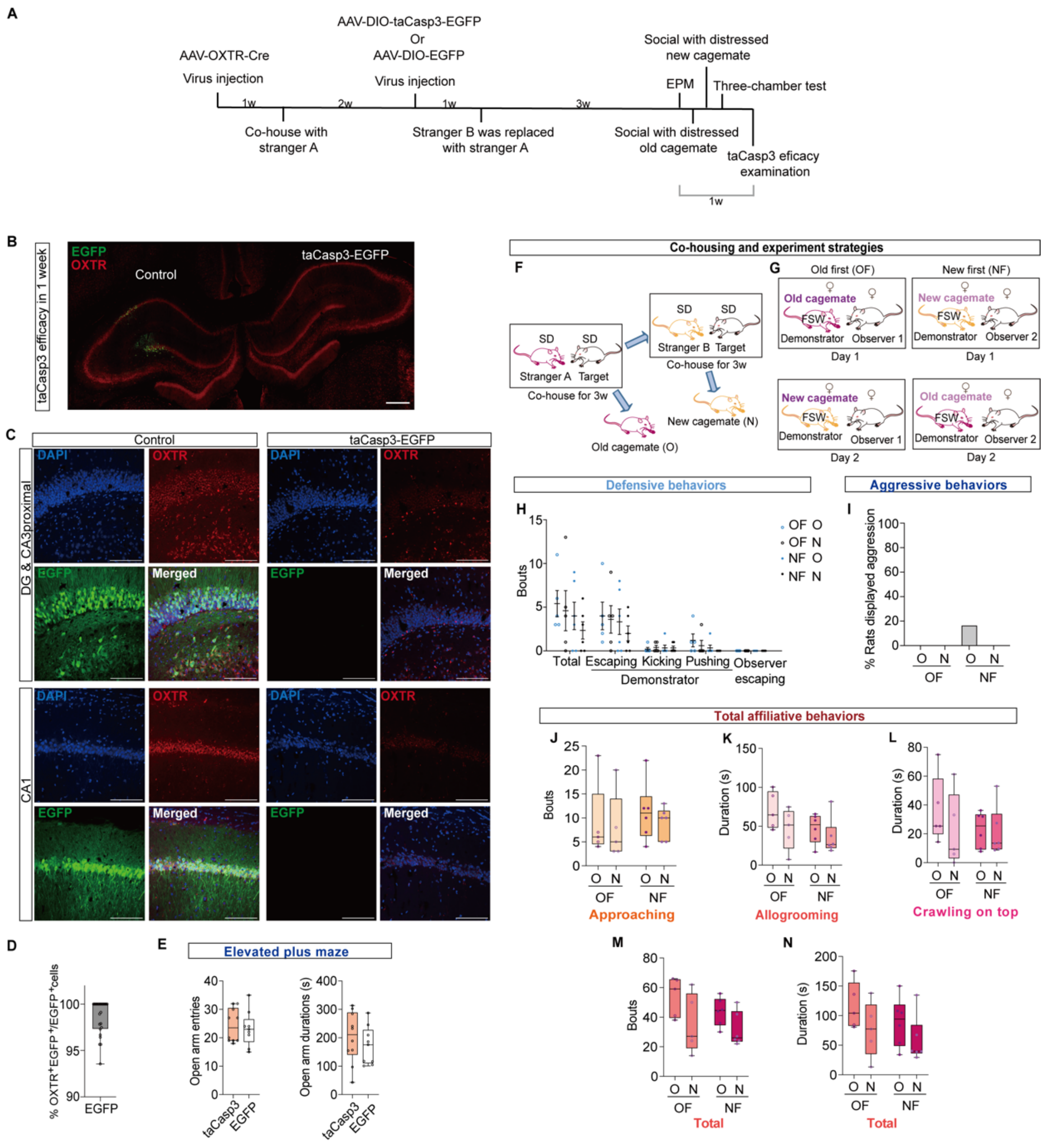

**Figure S3. Dorsal hippocampal OXTR-expressing neurons are necessary for adult rat peer bonding establishment.**

(**A**) Strategies for virus injection, co-housing and behavioral tests. (**B**) Representative images of OXTR expressions in rat after virus were injected for 1 week. Scale bar, 500 µm. (**C**) Representative images of OXTR expressions in dorsal hippocampus (DG, $CA3_{proximal}$ and CA1) of treatment (OXTR::taCasp3-EGFP, right) rats and control (OXTR::EGFP, left) rats after virus were injected for 5 weeks. Scale bar, 100 µm. (**D**) Ratio of $OXTR^{+}$ cells in $EGFP^{+}$ cells in control rats. $N_{total}$=24 (8 rats and 3 samples for each rat). (**E**) Comparison of behaviors in elevated plus maze test. n=9-10, Mann-Whitney test (left) or unpaired Student's *t*-test (right), no significance. (**F**) Co-housing strategy. Stranger rats that co-housed for the first 3 weeks were old cagemates, while those co-housed for the another following 3 weeks were new cagemates. (**G**) Experimental setup. On the first experimental day, observer rats would be divided into two groups randomly. They would social either with old (old first) or new cagemate rats (new first) that suffered FSW. On the second day, those had social interaction with old cagemate on the first day would social with new cagemate and vice versa. (**H**) Comparison of defensive behaviors in old first group (OF) and new first group (NF).

$n_{old\ first}$=5, $n_{new\ first}$=6, Two-way repeated ANOVA followed by Šídák's multiple comparisons test, no significance. (**I**) Ratio of aggressive behaviors in each group. (**J-N**) Comparison of affiliative behaviors in old first group (OF) and new first group (NF). $n_{old\ first}$=5, $n_{new\ first}$=6, Two-way repeated ANOVA followed by Šídák's multiple comparisons test, no significance.

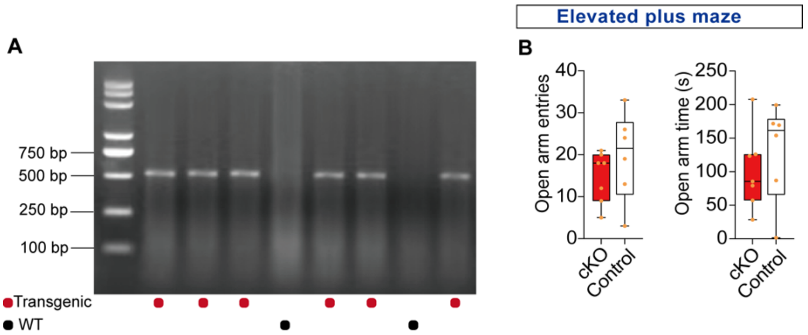


**Figure S4. *Oxtr* regulates peer bonding establishment in dorsal hippocampus.**

(**A**) Genotyping of LSL-Cas9 rat. (**B**) Comparison of behaviors in elevated plus maze test. $n_{cKO}$=7, $n_{control}$=6, unpaired Student's *t*-test, no significance.

**Movie S1. Agonistic and affiliative behavior types.**

**Movie S2. Social behaviors between distressed peer adult rats.**
